# High changing curvature regions detect chromatin enrichment in single cell data

**DOI:** 10.1101/2023.03.31.535120

**Authors:** Giulia Amici, Andrea Papale, David Lando, Wayne Boucher, D. Holcman

## Abstract

Chromatin organization is nowadays accessible from population and single cell Hi-C data in the form of large contact matrices. Yet reconstructing the three-dimensional chromatin structure remains challenging and relies on polymer models and numerical simulations to account for these Hi-C data. Here we propose a novel optimization algorithm to identify cross-linker ensemble reproducing the experimental contact matrix. Furthermore, based on the polymer configurations extracted from the optimization procedure, we identify high changing curvature (HCC) regions in the chromatin, detected both in population and in single cell Hi-C, that we also compare to Topological Associated Domains (TADs). We report here that the HCC regions correlate with gene expression and CTCF high density distribution. Interestingly, the HCC region locations are heterogeneous across one cell repeats, revealing cell-to-cell variability. To conclude, HCC regions appear both in single and population Hi-C polymer reconstruction and can provide a possible unit for gene regulation.

## INTRODUCTION

Population Hi-C experiments in eukaryotic cells[1, 2] have revealed a multi-layered organization of chromatin [3], from Topological Associated Domains (TADs) to chromosome territories [4, 5], and also to reconstruct tri-dimensional arrangements [6]. The combination of Hi-C data and polymer models [7–12] allows one to reconstruct chromatin organization at various spatial resolutions and thus to investigate the correlation between the structural organization with gene expression and regulation [13–17].

However, population Hi-C data give aggregated information collected from millions of cells, whereas information concerning a single structure only became accessible after the introduction of the single-nucleus Hi-C experiments [12, 18–22], sampling contacts of one single cell at a time. Nevertheless, structural organizations, such as TADs, emerge from population average sampling of many thousands of pairwise interactions, or loops between two genomic loci (for a recent review see Hafner and Boettiger [23]). Due to the heterogeneity of single cells only a very small subset of these pairwise interactions, or loops may exist in any given cell. It thus remains unclear how to quantify and study the chromosomal structural organization of TADs at a single cell level. For example, the chromatin topology that describes the ensemble of loops, their curvatures, torsions, proximity in three dimensional space with other loops remains difficult to quantify for several reasons: first loops can appear and disappear constantly [24, 25], second, although the notion of proximity is well captured by the experiments involving cross-linkers, it remains more difficult to quantify these distance using theoretical tools. To study chromatin organization, we propose to quantify the chromatin topological properties by computing the curvature, the torsion along the genomic length a procedure that reveals highly curved portion. We will correlate these properties with population organization such as TADs and also withCTCF bindings and gene expressions. Indeed, curvature analysis was previously used to quantify the shape of proteins [26–29] and we shall use these properties to locate sequences with high changing curvature (HCC) regions from mouse chromosomes. To find such regions we introduce an ensemble of computational steps starting from contact probability matrices, obtained either from single or population Hi-C. Then, we introduce an iteration procedure based on attributing cross-linkers to a polymer model [30–32] and molecular dynamics simulations, to find the optimal coarse-grained representation for chromatin, comparing simulated and experimental Hi-C contact matrices.

## MATERIALS AND METHODS

### Experimental data description

#### Hi-C experiments

Single-nucleus and population Hi-C processing and library preparation were carried out as previously described [18] using a haploid mouse embryonic stem cell line [33]. Libraries were sequenced on an Illumina HiSeq platform with 2×125bp paired end reads. Details of the samples used in this study can be found in Data Table I. The sequencing libraries were processed using our software package “NucProcess”, available at: https://github.com/TheLaueLab/nuc_processing.

**Table I.**
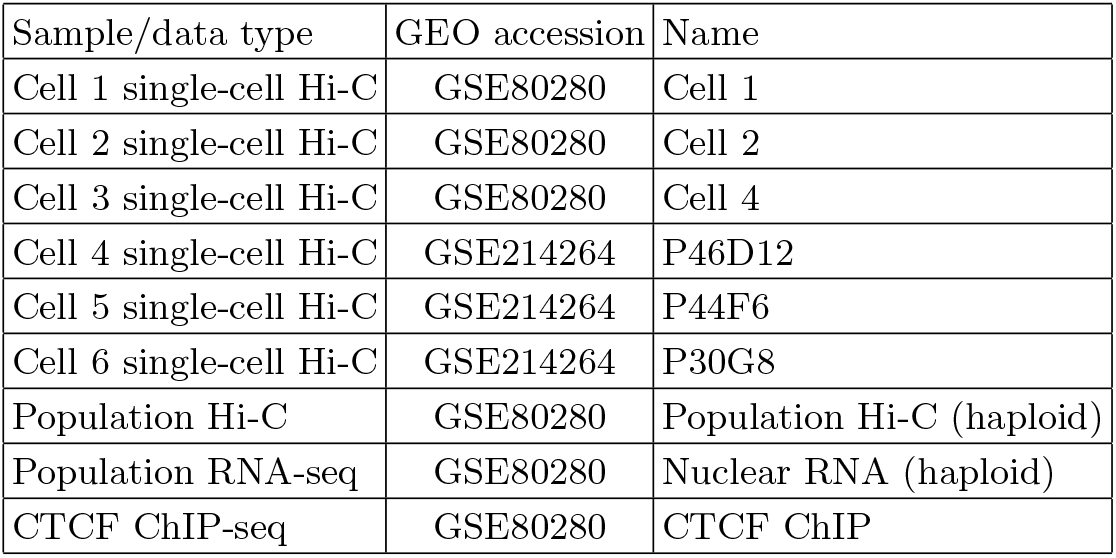
Details of the datasets used in this study deposited at Gene Expression Omnibus repository.

**Table II.** List of parameters used in the numerical simulations.

| Parameters | Symbols | Values |
| --- | --- | --- |
| Number of beads | N | 2334 (chr 19)<br>3677 (chr 17) |
| Number of cross-linkers | $N_c$ | 1282 (chr 19)<br>2029 (chr 17) |
| Friction | $\gamma$ | $0.5\tau_{LJ}^{-1}$ [35] |
| Energy scale | $\epsilon = \kappa_B T$ | 1 [dimensionless unit] |
| Mass | $m$ | 1 [dimensionless unit] |
| Monomer size | $\sigma$ | 1, [25kbp] |
| Integration time step | $dt$ | $0.012 \tau_{MD}$ |
| Elastic constant | $k_{rc}$ | $0.1 \epsilon$ |
| Maximal bond extension | $R_0$ | $1.5 \sigma$ |
| Persistence length | $l_p$ | $0.5 \sigma$ |
| Contact parameter | $d_{cut}$ | $2.5 \sigma$ |

Briefly, this software takes paired FASTQ sequence read files, a reference genome sequence and knowledge of experimental parameters (restriction enzyme type and the range of size of DNA fragments sequenced in the library) to create processed, single-cell or population contact files [12, 18].

#### RNA-seq experiments

RNA-seq samples were prepared for sequencing using a haploid mouse embryonic stem cell line as previously described by Stevens et al 2017 [18], and the data deposited at Gene Expression Omnibus, accession number GSE80280-see Data Table I. Briefly, the analysis of transcript abundance estimates for all annotated mouse transcripts (Ensembl v71, GRCm38/mm10) and ERCC spike-in sequences were obtained using Kallisto v0.42.4 [34].

#### CTCF-peaks experiments

CTCF chromatin immunoprecipitation (ChIP) sample from haploid mouse embryonic stem cell line was prepared and processed as previously described by Stevens et al 2017 [18], and the data deposited at Gene Expression Omnibus, accession number GSE80280-see Data Table I. Briefly, fixation was carried out using 1% formaldehyde for 10 minutes at room temperature and quenched with 150 mM glycine. DNA was fragmented in the presence of 1.0% SDS using a Bioruptor sonication instrument (Diagenode) producing a size range of 200 to 300 bp. The libraries were prepared using the NEXTflex Rapid DNA-seq kit (Illumina) and sequenced on a Hiseq-4000 platform with 2 × 50 bp paired end reads. Paired-end reads were trimmed with Trim Galore v0.4.0 using default parameters prior to mapping to the mouse genome reference sequence (GRCm38.p6) using Bowtie 2 v2.1.0. Peaks were called using MACS2 v2.1. using the narrowPeaks mode. Finally, read abundance profiles for each CTCF peak were calculated from the normalised bigwig files using Deeptools.

### Polymer model for chromatin structure

Chromatin is represented as a semi-flexible chain of *N* beads [35, 36], with diameter *σ* corresponding to 25kbp. The dynamical evolution of each bead of the polymer chain follows Langevin’s equation:

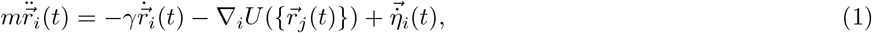

Where 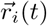 is the position of bead *i* at time *t* and *U* is the potential applied on bead *i* which depends on the positions of the overall polymer, *m* is the bead mass, *γ* is the friction coefficient and *η* is a Gaussian noise acting on each bead *i. η* has zero mean and is delta-correlated, satisfying the fluctuation-dissipation relation:

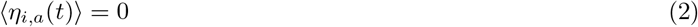

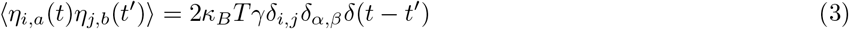

where *κ*_*B*_ is the Boltzmann constant, *α, β* and *i, j* represent the Cartesian coordinates and the particle indices respectively. *δ*_*i,j*_ is the Kronecher delta function and *δ*(*t* − *t*′) is the Dirac delta. The potential energy defining the polymer dynamics in eq. 1 is given by the sum of different energies:

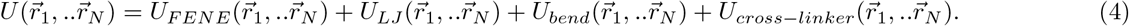

We shall now explicit each of them.

1. **The finitely extensible nonlinear potential (FENE)** bounds consecutive monomers along the linear chain following the expression

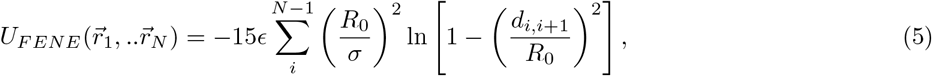

where the bond length 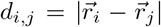 (distance between two monomers) cannot be larger than the constant *R*_0_ = 1.5*σ*. Here *ϵ* = *k*_*B*_*T* is the energy.
2. **Truncated and shifted Lennard-Jones potential** is applied on all monomers following short range excluded volume interactions and the associated energy is given by

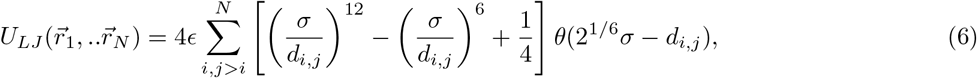

where θ is the Heaviside function.
3. **Bending energy** 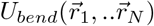 penalizes consecutive bond vectors 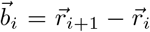 that are not parallel. The chain rigidity is enforced by the following expression for the energy

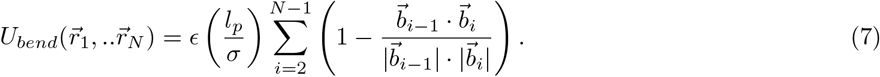

Since the bead diameter corresponds to a length *σ* = 25kbps, the persistence length *l*_*p*_ of the chain is chosen equal to *l*_*p*_ = 0.5*σ* [9, 11].
4. **Harmonic potential** *U*_*cross−linker*_ allows to add a force between N_*c*_ cross-linkers randomly chosen connecting non-consecutive beads [30, 31, 37, 38]. This term accounts for the presence of long-range interactions and reproduces the multi-scale organization emerging from Hi-C data. The energy is given by

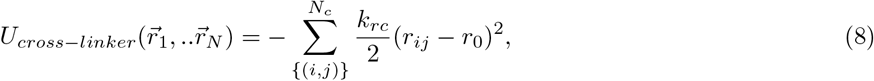

where the spring constant is given by *k*_*rc*_ = 0.1*ϵ*.

At the scale *σ* = 25kbp, the presence of fixed cross-linkers allow us to reconstruct chromatin structure by modeling the effects of cohesin and CTCF [32, 39] without including explicitly the extruding process occurring at a time scale of 1*kb*/s, which is much faster than the time scale we are interested in here [40–42].

#### Polymer stochastic simulation procedure

The numerical simulations are performed using fixed-volume and constant-temperature molecular dynamics (MD) with implicit solvent using the LAMMPS simulation package [43]. The Langevin equation 1 is integrated numerically using Verlet’s algorithm with the integration timestep sets to *dt* = 0.012*τ*_*LJ*_ and the local friction 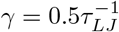, where τ_*LJ*_ is the characteristic simulation time given by 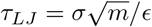 in Lennard-Jones units. We set the mass m = 1, the length *σ* = 1 and the energy scale *ϵ* = *k*_*b*_*T* = 1 in the dimensionless units. The polymer model is contained within a simulation cubic box with periodic boundary conditions large enough to avoid interactions with its own copy.

Each MD-simulation includes a short initial run with 10^4^ integration steps with a restrained dynamics (implemented using the *nve/limit* integration algorithm [43]) and with a reduced elastic constant 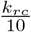. This choice allows to smooth the converging steps toward a final polymer configuration with the newly added random cross-linkers. Then we relax this condition and use the elastic constant *k*_*rc*_ to perform 2 · 10^6^ integration steps to reach equilibrium.

### Cross-linker selection for population versus single cell Hi-C

We shall explore here population and single cell Hi-C data with our simulations. In particular, we will use Hi-C data for the mouse chromosome 17 and 19 at 25kbp resolution. In the population Hi-C, every bead pairs is associated to a contact frequency, measured from a sample of ∼ 10^6^ cells. Using the Hi-C matrix as input, we will present below an optimization algorithm to estimate the probability for two beads to be connected by a cross-linker. Thus leading to a polymer model that best account for the local chromatin organization contained in the Hi-C matrix. From that probability distribution, we shall extract a set of *N*_*c*_ pairs that are cross-linked. Following this procedure, we will simulate 500 polymer realizations to account for the population heterogeneity.

For single cell Hi-C, six distinct single-nucleus Hi-C datasets have been used as an input to select the cross-linker location. In this case, we added a cross-linker to connect a pair of beads with an entry equal to 1 in the corresponding Hi-C matrix.

### The RCL-reconstruction (RCR) Algorithm to select cross-linkers from Hi-C population data

We describe now an algorithmic procedure to select a certain ensemble of connectors to define the chromatin local topology whose simulated Hi-C map is the best approximation of the experimental one. Using the Hi-C matrix as an input and, through *k* iterations, the algorithm provide a series of probability 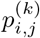 for each pair of beads *i, j* to be cross-linked in the chromosome model. Then we use the harmonic potential *U*_*cross−linker*_ to defines the bond energy between the set of *N*_*c*_ beads.

#### Initial step

At the initial step *k* = 0, we start with a null probability distribution 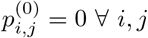 for which we ran 500 polymer realizations using self-avoiding chains. For each realization of cross-linkers, we sampled 50 independent configurations at equilibrium to compute the contact matrix *M*^(0)^ (Fig. 2A). This matrix is obtained by computing the frequency of interaction between each pair of monomers located at a distance smaller than *d*_*cut*_ = 2.5*σ*. Then, we compute the probability of having a cross-linker between *i* and *j* from:

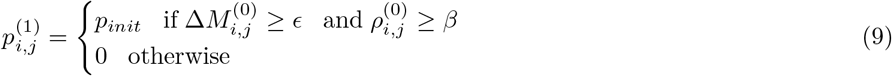

where we chose *p*_*init*_ = 0.01 and *β* = 0.5. The difference of Hi-C and simulated contact matrix at step 0 is defined for *i, j* by

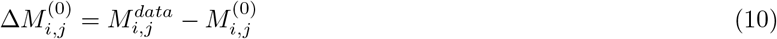

and the normalized difference is given by

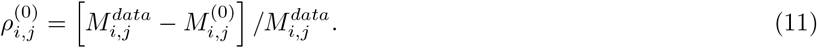

The threshold *ϵ* is defined by

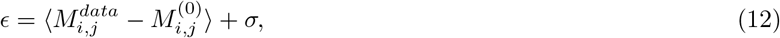

where the standard deviation 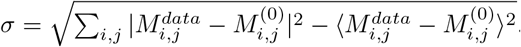. Note that when 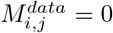, connectors cannot be added. The threshold *ϵ* has been introduced to update the probability of pairs of monomers only when the difference between simulations and data is such that the presence of a cross-linker is justified and not due to numerical errors or finite size effects. Then, we repeated this procedure using formula 9 and extracting 500 polymer realizations with 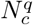 cross-linkers, where index *q* refers to the single realization.Then, using the final configurations of step 0 as initial states we performed MD simulations and computed the normalized contact probability matrix *M*^(1)^.

#### Iterative step

The procedure has then been iterated for k iterations (see Fig. 2B),

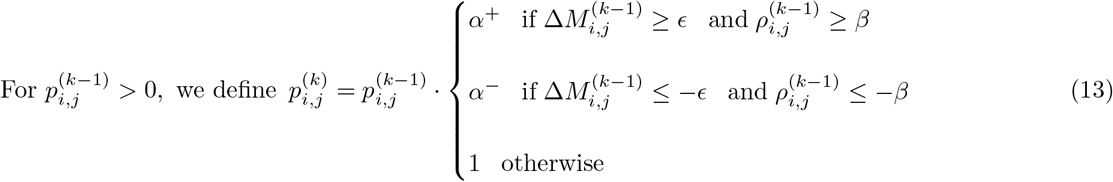

and

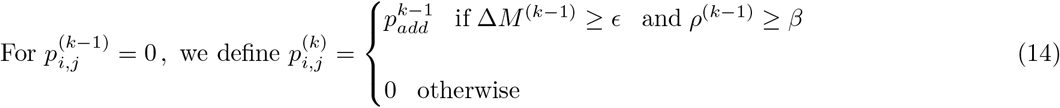

where

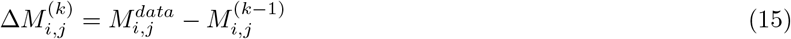

and the normalized difference is given by

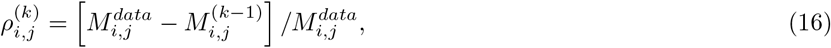

where 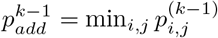. We set *α*^+^ = 1.5 and *α*^*−*^ = 0.5 to increase or reduce the probabilities recursively.

To evaluate the convergence of the probability series, we computed the stratum-adjusted correlation coefficient *SCC* [44] between the simulated and the experimental contact matrices, computed as the weighted average of the Pearson correlation coefficients for each sub-diagonal:

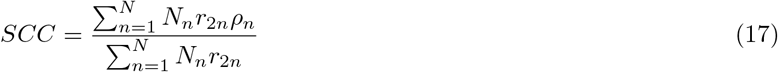

where the index *n* refers to the *n*−subdiagonal containing *N*_*n*_ elements, 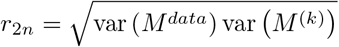 and *ρ*_*n*_ is the Pearson correlation coefficient. We also estimated the square error between the contact matrix computed at iteration *k* and the experimental one:

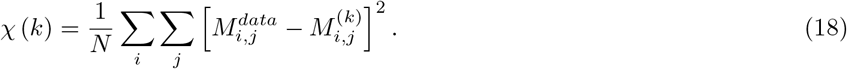

#### Termination step

The optimal contact probability is selected such as the stratum-adjusted correlation series coefficients *SCC*_*k*_ and the error function series χ (k) reach a steady-state. In practice, we used *k*_*final*_ = 15 (Fig. 2D). In that case, we chose to represent chromatin using the cross-linker probability:

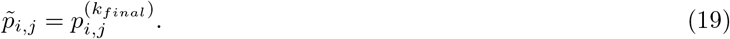

### Spline interpolation and High Changing Curvature (HCC) region detection

The reconstructed polymer configuration consists of a collection of discreet beads (Fig. 1B-a). However, to identify regions of high curvature, we decided to map the three dimensional discreet polymer representation into a continuous one. For that goal, we interpolated the cartesian coordinates 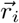 of each bead *i* = 1, .., *N* by order three polynomial splines (Fig. 1B-b yellow) since we will need a continuous curve up to its third derivative in the following analysis.

**Figure 1.**
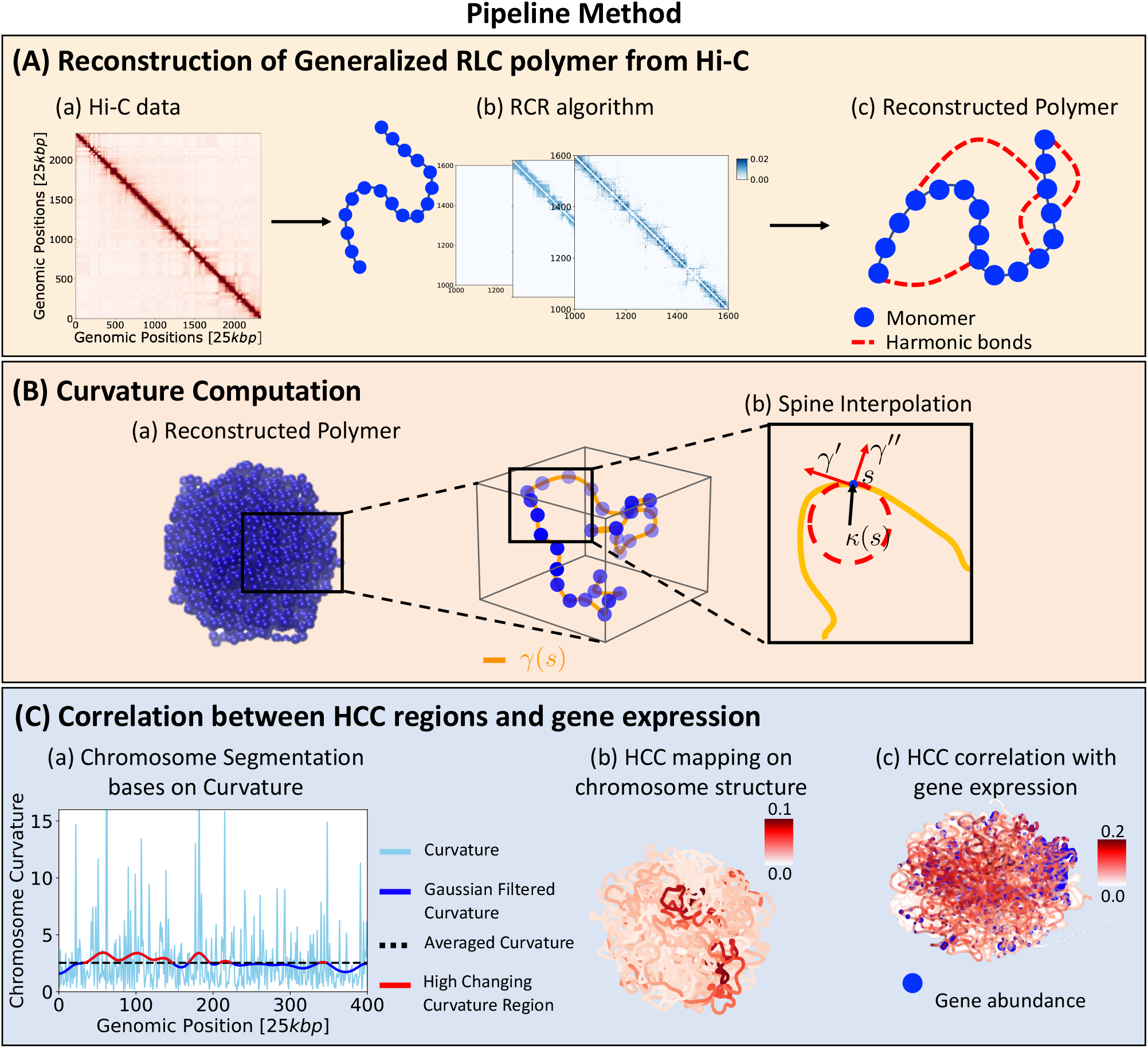
Schematic of the framework to detect High Changing Curvature regions. Panel A. Using population Hi-C matrix for mouse chromosome 17 and 19 (sub-panel a) at 25 *kbp* scale, we developed a recursive optimization process (RCR, sub-panel b) based on a generalization of the randomly cross-linked model (c) to reconstruct the chromosome spatial organization. Panel B. The simulated polymer structure (sub-panel a), as discrete curve, can be numerically interpolated using spline interpolation method to compute a smooth curve (sub-panel b). Panel C. We have computed the curvature of the smooth curve (light blue line in sub-panel a) as a function of the genomic position and its filtered curvature (blue line) using a Gaussian kernel with standard deviation equal to 10. Thus, we define High Changing Curvature regions (red segments) the sequences where the curvature is larger than its average value (black bashed line). Then, the probability of having an HCC region is highlighted (sub-panel b) for a reconstructed configuration. Finally, the HCC position distribution averaged over the ensemble of polymer realizations is compared with gene abundance distribution on the same chromosome (sub-panel c).

**Figure 2.**
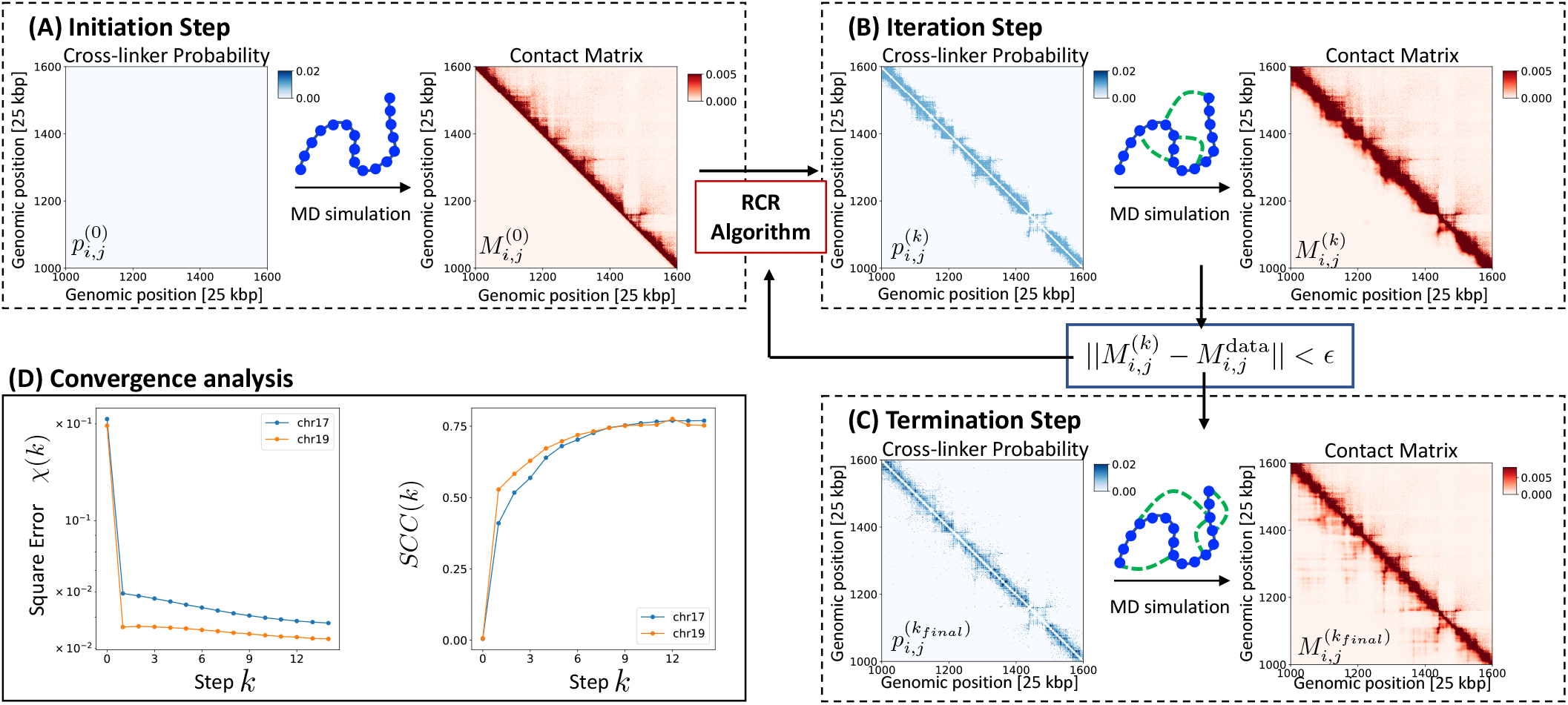
RCL reconstruction (RCR) Algorithm. The ensemble of cross-linker to reconstruct numerically the optimal contact matrix is obtain using the RCR recursive algorithm for *k* iterations: (A) Initiation step *k* = 0: MD simulation of 500 polymer configuration are performed without cross-linker 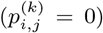 to compute *M* ^(0)^. (B) Iteration step *k*: applying eqs. 13-14 to the result of the previous iteration, the new cross-linker probability 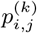 is computed, 500 polymer realization are extracted and their MD simulations performed to compute *M* ^(*k*)^. (C) Termination step *k*_*final*_: when the error function reaches a steady-state the iteration is terminated. (D) Convergence: square error and stratum-adjusted correlation coefficient (SCC) as a function of the algorithm iteration step for chromosomes 17 and 19.

The spline interpolation procedure [26–29] is detailed in SI section.

The output of this procedure is smooth curve *γ* parameterized by the arclength *s* from which we can compute the curvature *κ*(*s*) and the torsion *τ*(*s*) according to the Frenet-Serret formulas[45]:

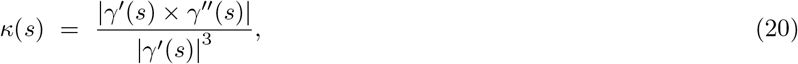

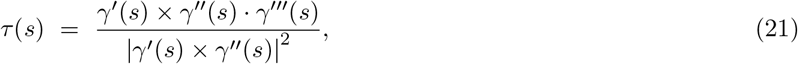

where the derivative is computed with respect to the arclength s. We recall that the geometry of the curve is fully described by the curvature *κ* and the torsion *τ*, but we decided to focus on computing the curvature to quantify the regions of high curvature inside the polymer configuration, that accounts for the chromatin organization.

To detect the subregions of chromosomes characterized by a high curvatures, we use the following criteria: first we computed *κ*(*s*), then we computed as a threshold the mean curvature *K* over the entire polymer (Fig. 1C-a):

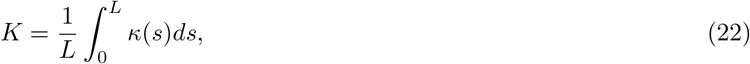

where *L* is the total length of the polymer interpolated the spline. Thus, the *High Changing Curvature regions* (HCC regions) are defined by

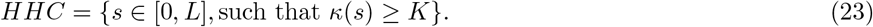

HCC consists of a union of genomic arclength intervals (Fig. 1C-a).

Holcma12

### Filtering the high fluctuating curvature by a Gaussian

Due to the high variability of the curvature along the genomic length, we convolve the curvature with a Gaussian filter

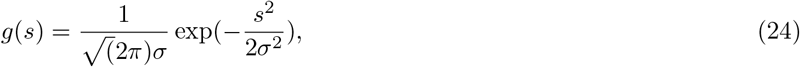

where the standard deviation *σ* is chosen 10. The output of this procedure is a smoother curvature at scale *σ*, defined by

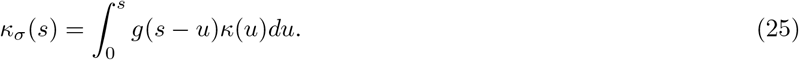

## RESULTS

The goal of the method we developed here is to localize HCC regions in the chromosome tridimensional structure. In order to do that, we started from Hi-C data for mouse at 25 *kbp* resolution, in both population and single cell. Using the RCR-algorithm (method section), we converted the Hi-C map into an ensemble of cross-linkers. We included these connectors to a semi-flexible polymer (method subsection) and then we simulated this polymer model to reconstruct chromatin structure (Fig. 1A). To convert the discrete polymer configuration in a continuous chain, we used spline interpolation to obtain a smooth chromatin curve (Fig. 1B). From this procedure, we computed the curvature along the genomic distance and detected the HCC regions along the chromatin (Fig. 1C-a). Finally, we will use this procedure to correlate the HCC regions with gene expression and CTCF peaks data (Fig. 1C-c).

### Population HiC

We first used the RCR-algorithm to construct a polymer model associated with the contact matrix of mice chromosomes 17 and 19. We compare the reconstructed matrix (Fig. 3A upper) from the algorithm with the experimental one (Fig. 3A lower) by computing the stratum-adjusted correlation coefficient [44], with a score of *scc* = 0.85. This high value assesses the reliability of the reconstructed polymer to reproduce the experimental data.

**Figure 3.**
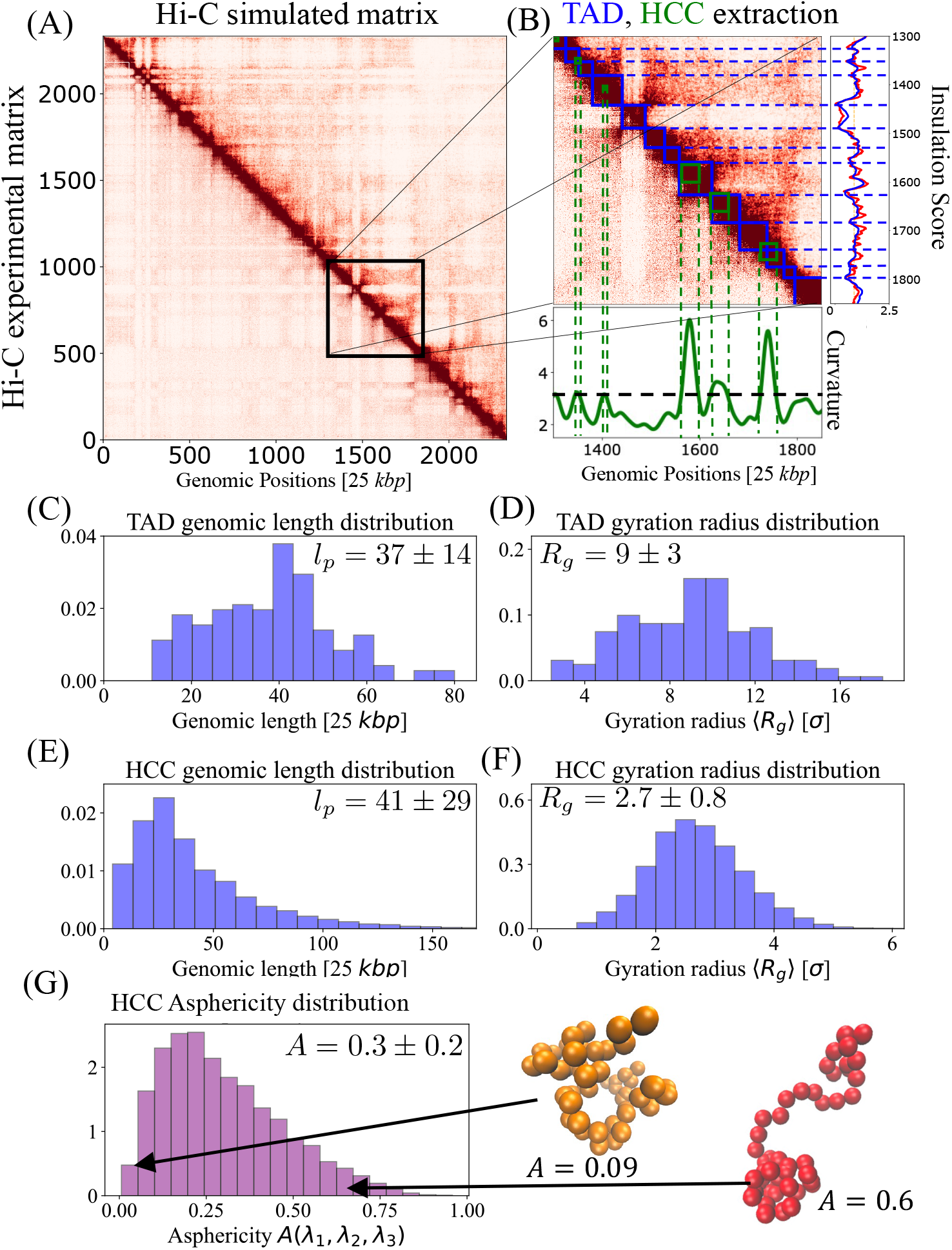
Results for population Hi-C dataset. (A) Simulated Hi-C frequency matrix for mouse chromosome 19 at 25 *kbp* resolution. (B) Zoomed section of the Hi-C matrix, where we focus on TADs, detected from the insulation score (experimental, red line, and simulated, blue), depicted as blue squares, and HCC, extracted from curvature (green line), as green squares. We report here genomic length and gyration radius distributions computed on the ensemble of 500 polymer realizations respectively for TADs (C-D) and HCC (E-F). (G) Finally we have analyzed the tridimensional shape of HCC, computing the asphericity distribution. Examples of HCC with high and low asphericity are shown.

To further compare the contact matrix computed from the polymer model with traditional HiC characteristics, we first located TADs on the simulated matrix, using the Insulation Score [46] (Fig.3B, blue squares). We found 56 TADs, with a mean genomic length ⟨*ℓ*_*p*_⟩ = 925 ± 400 kbp, (see also the full distribution in Fig. 3C). Moreover, we estimated the tridimensional size using the average TAD gyration radius

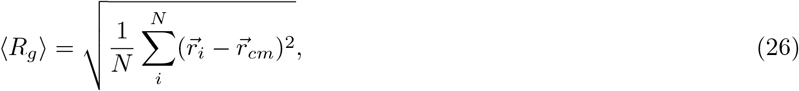

where 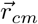 is th TAD centre of mass and the distribution is shown in Fig.3D. We found that the average gyration radius is ⟨R_*g*_⟩ = 9 ±3*σ*.

After we located TADs on the contact matrix, we investigated how they correlate with HCC regions. While TADs emerge as a *average* computed over many chromosome arrangements, HCC regions are associated with a single chromosome structure. Therefore, we computed the curvature profile for each realization that we average over the polymer realizations. We use the mean curvature criteria (method section) to identify the HCC region, as shown in Fig.3B (green squares). On average, we found an average of 39.8 HCC regions per polymer realization for chromosome 17 and 27.8 for chromosome 19. The genomic size and the gyration radius distributions of HCC regions are shown in Fig. 3E-F where the mean genomic length is ⟨*ℓ*_*p*_⟩ = 1025±725 kbp and the average gyration radius is ⟨R_*g*_⟩ = 2.7±0.8*σ*. To further compare TADs with HCC length, we estimated the ratio ⟨*ℓ*_*tad*_⟩/⟨*ℓ*_*hcc*_⟩ = 0.9, while the radius of gyration ratio is given by ⟨*Rg*_*tad*_⟩ / ⟨*Rg*_*hcc*_⟩ = 3.3. Interestingly, the HCC regions are *ρ*_*hcc*_/*ρ*_*tad*_ = 41 times denser than TADs (where the density ρ is computed as ∼ *ℓ*/*Rg*^3^). We also found that 44% of the TADs have at least a partial overlap with HCC, for a fraction of their genomic length corresponding to 57%.

There are two main preliminary questions about HCC regions. First: *where* are they situated along the chromosome? Second: *how* they are made? Do they have a peculiar shape or not?

To answer the first question, we divided the chromosome structures in three parts with the same length (begin, center,end) and we found that, averaging over 500 polymer realizations, HCC are localized for the 33.8%, 32.8% and 33.4% in each part of chromosome 17 and 31.9%, 33.7% and 34.4% for 19. This result suggests that HCC regions span the chromosome in its entire length. Moreover, on average they cover the 44% of the chromosome length (computed as ∑*HCC*_*i*_/ total chromosome length).

To gain insight of the tridimensional shape of these HCC regions, we computed the eigenvalues of the gyration radius tensor [47, 48]

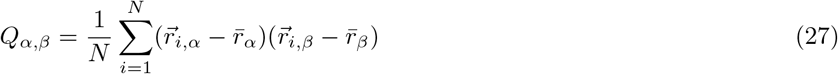

where *α, β* are the cartesian coordinates (*x, y, z*) and 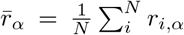 and *r*_*i,α*_ is the monomer i position. The symmetric 3×3 matrix *Q* has eigenvalues *λ*_1_, *λ*_2_, *λ*_3_, corresponding to the axes of the principal moments of inertia of the polymer. The shape of the polymer conformation can be quantified by considering the ratios of eigenvalues using the asphericity, that characterizes a deviation from a spherical symmetry:

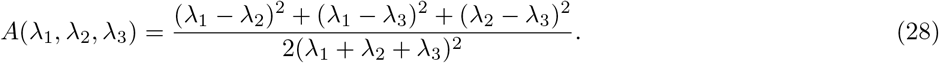

The asphericity *A*(*λ*_1_, *λ*_2_, *λ*_3_) varies from 0 to 1. When *A* = 0, *λ*_1_ = *λ*_2_ = *λ*_3_ and the polymer has a spherical shape. However, when *A* = 1, then *λ*_2_ = *λ*_3_ = 0 and the polymer has a rod-like shape. We computed the asphericity for the 500 simulated realizations of chromosomes 17 and 19 and found that HCC regions are heterogeneous, as shown by the distribution of Fig.3G.

### HCC regions correlate with gene expression

After we segmented HCC regions, we investigated whether they could have a functional role so that high curvature regions correlate with gene expression. Gene positioning in the nucleus is often related to the transcriptional activity [13–17], that could be influenced by the local organization and chromatin condensation. We first estimated the occurrence of having a HCC region on a given genomic position (Fig.4A red intensity) that can be compared with the level of gene abundance (blue circles). We quantify their correlation using the Spearman coefficient [49] and found *S*_*sper*_ = 0.293 and *S*_*sper*_ = 0.458, as shown in Fig. 4B for chromosome 17 (left)and 19 (right), suggesting that the presence of high curved chromatin is slightly correlated with the presence of expressed genes. This result is in contrast with the case where there are no random cross-linkers (*N*_*c*_ = 0), where we found that *S*_*sper*_ = − 0.05, after we ran 500 polymer realizations. Here, the simulated contact matrix does not account for the experimental Hi-C matrix and the chromatin structure is not reproduced correctly.

**Figure 4.**
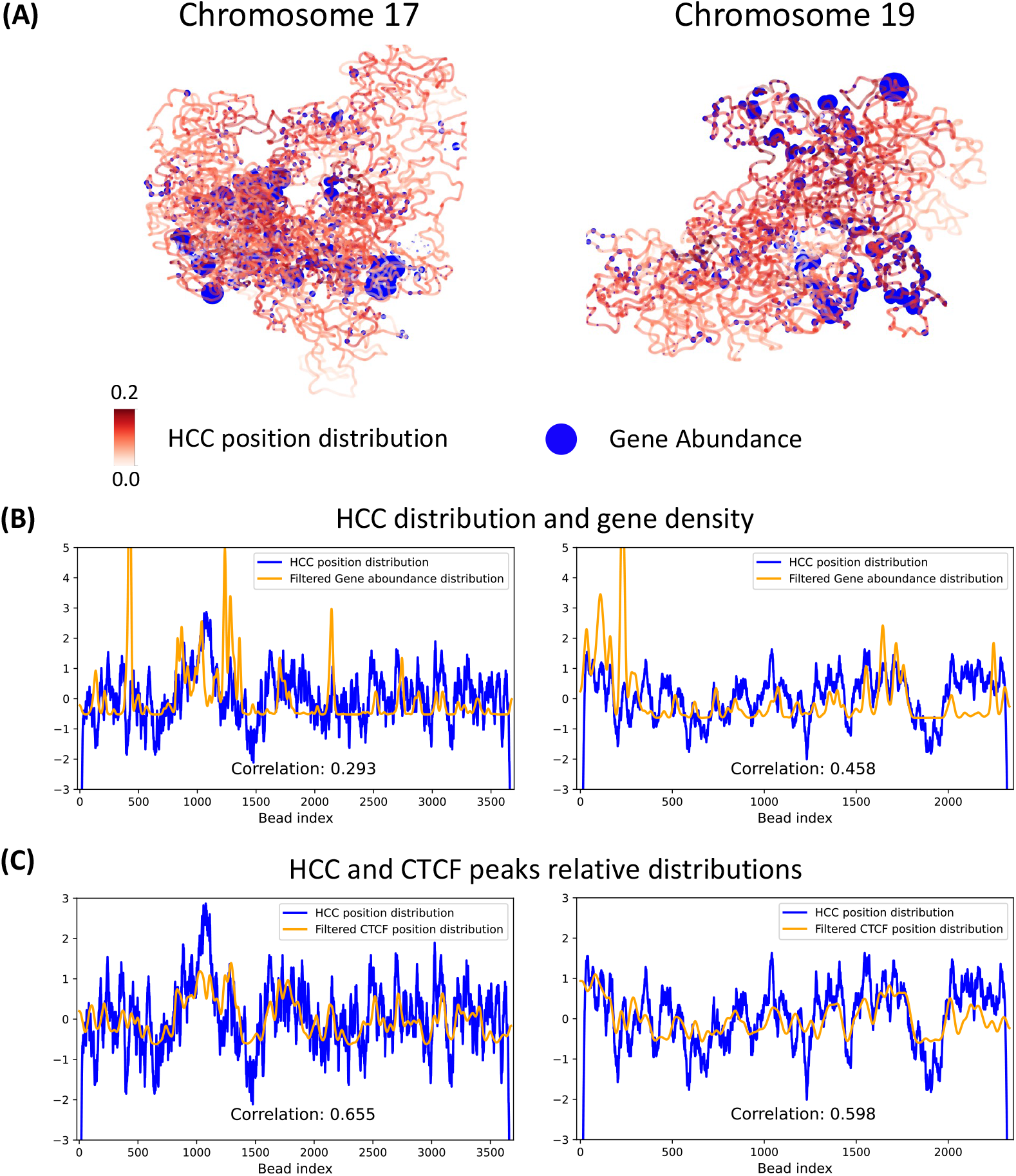
(A) Polymer structures colored according the averaged distribution of HCC regions alongside with blue spheres representing the presence of genes whose size depends on their abundance for chromosome 17 (left) and 19 (right). (B-C) Comparison between the averaged distribution of HCC regions on the chromosome 17 (B) and 19 (E) with the experimental gene abundance distribution, as a function of the genomic position. To compare two distributions computed at different scales we applied a Gaussian filter with standard deviation *σ* = 10 on the gene abundance distribution. (D-E) Comparison between the averaged distribution of HCC regions on the chromosome 17 (D) and 19 (E) with the CTCF binding site distribution, as a function of the genomic position. To compare two distributions computed at different scales we applied a Gaussian filter with standard deviation *σ* = 10 on the gene abundance distribution

We also investigated the relation between the CTFC peaks distributions and the HCC regions (Fig. 4c) where we found that the Spearman correlation coefficient is given by *S*_*sper*_ = 0.655 (chromosome 17) and *S*_*sper*_ = 0.598 (chromosome 19). To conclude, these results show a relative correlation between the HCC, gene expression and CTCF distributions. Possibly, CTCF is involved in generating high curvature regions.

### HCC regions hidden in single cell Hi-C

Single cell Hi-C dataset is used to access the chromosomal structure a specific moment of a single cell cycle. Thus we decided to investigate how the cell-to-cell variability can influence chromosomal arrangement and in particular how the HCC region could vary, despite the fact that only 2.5% of all interactions are captured [19–21]. To identify the HCC regions, we started with the GRCL-polymer model. Due to the sparsity of the experimental *contact* matrix for Cell 1 (Fig. 5A upper matrix)), we added cross-linkers each time there was a non-zero probability in the contact matrix. We then simulated six different chromosome 19 starting from 6 single-cell Hi-C datasets and computed the simulated contact matrix (Fig. 5A lower matrix)).

**Figure 5.**
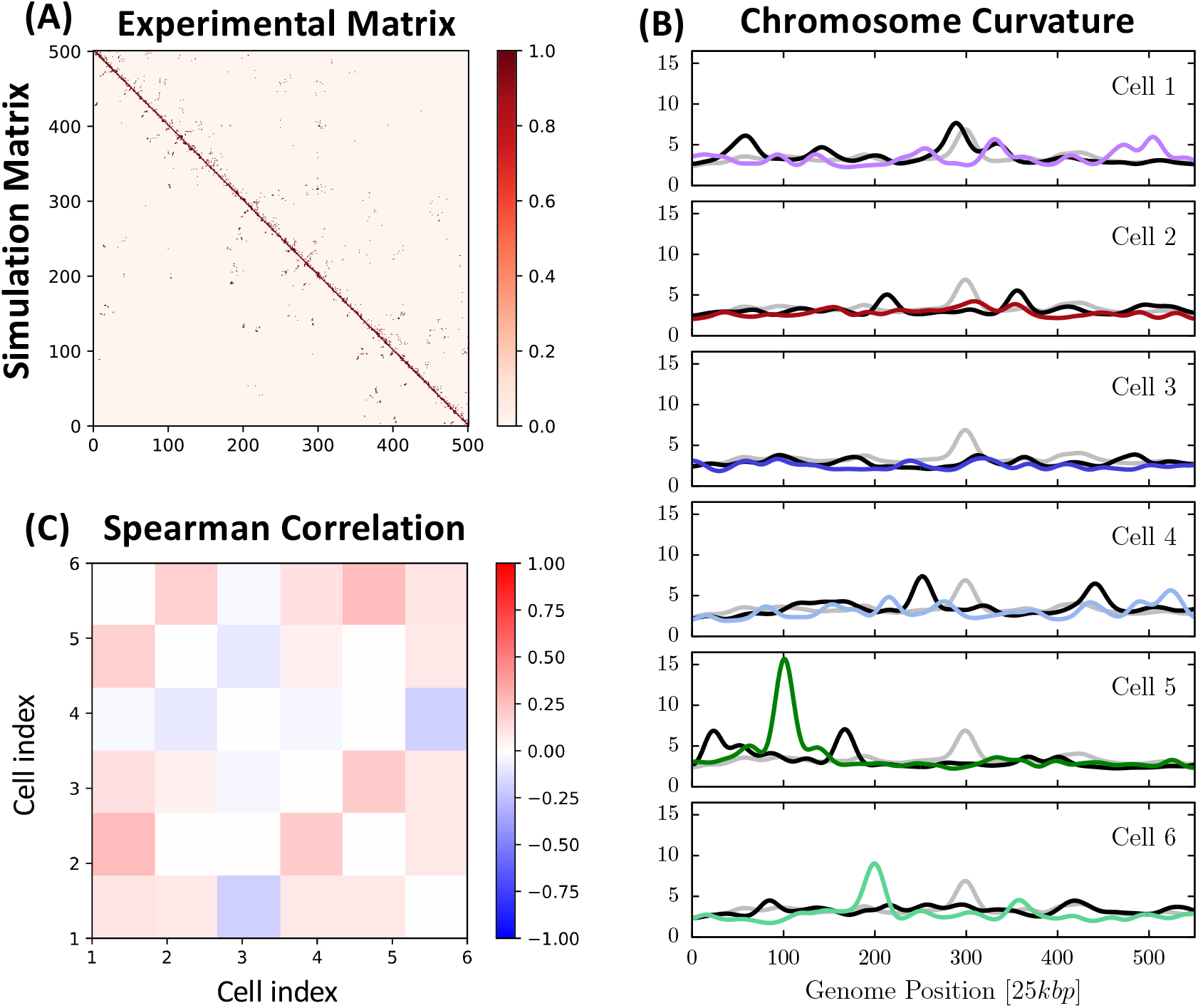
(A) Comparison between experimental Hi-C matrix (upper matrix) and simulated matrix (bottom matrix) for Cell (B) Curvature profile for the 6 simulated realizations of chromosome 19, as a function of the chain beads (colored lines). Grey lines: curvature averaged over all the 6 cell curvature profiles (colored line in each graph). Black likes: average curvature for 10 repetitions of the chromosome simulation at fixed cell. (C) Heat map of Spearman correlation among all possible pairs of cells.

To identify the HCC region, we applied the spline interpolation procedure (subsection of Method)on the 6 simulated chromosomes and finally obtained the curvature vs the genomic distance (Fig. 5B, colored lines: purple, brown, blue, green). We further plotted on each subfigure the single cell average (black) and the average over the six realizations (grey). We identify the HCC regions using the criteria that the curvature is above the mean (eq. 23) and found that on average (over six cells), there are 20 HCC regions in a chromosome, while the average HCC genomic length is ⟨*ℓ*_*p*_⟩ ∼ 1.1*Mbp* and average gyration radius ⟨R_*g*_⟩ ∼ 3.68*σ*.

Similar to the Hi-C population, HCC regions are not located in any preferential position with roughly %33.0, %38.1 and %28.9 occupancy at the beginning, the centre and at the end of the chromosomes). Finally, we calculated the Spearman correlation coefficient *S*_*sperm*_ of the curvature *κ*_*i*_(*s*_*q*_) of cell *i* at genomic distance *s*_*q*_ with the same for cell *j*, leading to the correlation matrix (Fig. 5C), showing a poor correlation between the curvature profiles. This result shows a large cell-to-cell variability associated with the chromatin reorganization over time.

## DISCUSSION

HCC regions are a generic tool for studying relevant 3D arrangements in the genome. In principle it could be used whatever is the experimental technique or the computational model behind the inferred chromosome structure. In particular, after applying it first to population Hi-C data, for which we explored the relationship of HCC sequences with TADs, we show that this tool can be applied to single cell Hi-C data, for which the usual definition of TADs cannot be used and the presence of stable genomic tridimensional arrangements has not been established yet. Second, we use gene expression abundance data and CTCF peak data in the population approach, and show that HCC region positioning shows correlation with the chromosomal gene abundance distribution and CTCF binding sites position distribution. Moreover we discover that, in the single cell approach, HCC positioning along the chromosome shows high cell to cell variability, indicating that HCC regions occurs differently for the same cell structure but captured in different single nucleus experiments. Taken together, these results suggest that HCC regions could have a functional role in gene expression and lay the foundation for a new point of view in studying the interplay between tridimensional folding and biological functionality.

The concept of curvature has been used to characterize protein folding [26, 50, 51] and to extract physical properties. Using differentiable density estimators such as curvature or torsion, it is possible to map protein structure into interactive structure. These tools are used to study homologous proteins that are relatively well conserved. Any amino acid substitutions can lead to differences in the structures. Curvature and torsion are used to classify fragments of the protein cores of homologues for an accurate comparison. Another application concern the representation of polypeptide chains as discrete curves to obtain a three-dimensional model from secondary structure [52]. However, much less results are known for quantifying Chromatin using curvatures due to the difficulty of reconstructing a smooth from a discrete polymer model.

### Algorithms for polymer reconstruction from Hi-C matrices

Over the past decades, polymer models have been systematically used to access chromatin statistics, once a criteria of reconstruction has chosen. Indeed, one of the main difficulty was to identify the polymer model that best represents the chromatin portion to be studied. This optimal reconstruction is obtained by comparing simulated versus experimental contact matrices. This construction and comparison involve the choice of a method, algorithms and parameters. Several computational procedure to reconstruct a bead-spring polymer model are possible [38], [53] in the absence of exact constraint. However, when constraint on a contact matrix, specific constraints are required: in a two steps algorithm, where the effect of repulsion forces between monomers is ignored [31], the method relies on choosing a minimal number of random connectors inside a restricted region. The polymer model accounts for diagonal blocks, while long-range specific genomic interactions were added based on the contact matrix.

The distribution of power value (called exponent *β*) values extracted from fitting the encounter probability (EP) by 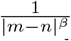 is the first step to recover the number of connectors, represented by springs between monomer-pairs and accounted for the chromatin architectures. Adding connectors between random monomers is sufficient to recover TAD sub-regions. Then it remains to add random and persistent long-range connections between monomers. Randomising connectors positions is sufficient to reproduce the inherent variability in chromatin architecture of nuclei population captured in 5C experiments. Accounting for long-range deterministic and short-range stochastic interactions leads to a more accurate polymer model for chromatin reconstruction. The quality of this approximation is measured by the decay norm (Kolmogorov-Smirnov distance) of the *β* – exponent between the data and the simulations. This approach can generate statistics of passage times and radius of gyration, which characterizes more accurately chromatin dynamics. The added connectors between random monomer-pairs characterizes sub-configurations present in 5C data.

This algorithm was generalized to reconstruct chromatin of mammalian nucleus and multiple Topologically Associating Domains (TADs) across cellular differentiation. To account for multiple interacting TADs, the parsimonious model accounted for randomly cross-linked (RCL) polymer model to map Hi-C encounters within and between TADs into direct loci interactions using cross-links at a given base-pair resolution. Such approach was used to study single particle trajectories, first passage times, predicting anomalous exponents and for studying synchronous compaction and decompaction of TADs throughout differentiation. Transient properties of the chromatin are inferred from steady-state statistics embedded in the Hi-C/5C data [54]. Recently, a Bayesian approach [25] with a similar spirit as [31] was used to reconstruct cross-linkers relying on a CTCF to model the local loop formation. A generalized Rouse Chromosome Model accounting for CTCF/cohein-mediated loops was previously used to compute the contact probability and the mean spatial distance [55]. Interestingly, mean distance matrices between loci measured at a single-cell level can be used as input to generate an ensemble of 3D polymer chromosome conformations [56]. The ensemble of configurations are obtained by maximum-entropy (exponential of the sum of all squared distances) criteria to recover higher-order structures such as three locus interactions, radius of gyration or clustered structures. Variant of polymer models are also used to generate formation of chromosome loops [57] or to account for the heterogeneity in chromosome organization.

Chromosome structure can also be reconstructed using long-range harmonic interactions among monomers [11]: this approach relies on a Gaussian polymer model, where the optimal coupling coefficients of the harmonic bonds are inferred from the experimental contact probabilities via an optimization procedure based on a gradient descent. The descent procedure is based on minimizing the energy landscape based on the distance between the Hi-C data and the contact probability matrix. The final result is a mean-field contact matrix that approximates the experimental Hi-C matrix. Similarly, the polymer-based recursive statistical inference method (PRISM) [55] uses strings and binders switch (SBS) polymer model [53], where long-range interactions among monomers are mediated by particles (binders) that can be bound at specific loci along the chain. The different blocks defining TADs in contact matrices are obtained by defining various groups of binding particles and by locating binding sites at appropriated positions. The correct binding sites and the number of binding particles are obtained by minimizing the distance between the input Hi-C matrix and the numerical contact matrix with a correction given by an additional Bayesian term to reduce overfitting. It has been applied to several experimental data (Hi/C, GAM, SPRITE) [58].

### TADS versus HCC

TADs represent a statistical averaging of chromatin organization derived from many millions of cell conformations and their cross-linkers. In contrast HCC regions are derived directly from the shape of the chromatin path and thus is well suited to studying these conformations in single cells. We report here that it is sufficient to have just tens of cross-linkers in a polymer chain of thousands of monomers to generate and detect high curvature. Thus a small change in the connectivity of cross-linkers can drastically influence the three dimensional folding and organization of chromatin. Currently, the loop extrusion mechanism is proposed to be the major force driving chromatin folding seen in Hi-C contact datasets [41, 59]. At the moment we do not understand the relationship we see in HCC regions and the loop-extrusion mechanism of chromatin folding. It could be that loop extrusion by the cohesin complex facilitates the formation of HCC regions or disrupts them. Further work analysing HCC in mutants involved in loop extrusion, such as SMC3 or WAPL, will be needed to address this question. As well, recent imaging studies have revealed that chromatin loops formed by loop extrusion are also highly dynamic [60], with the proteins involved also forming rapidly exchanging complexes [61]. Therefore, the high frequency at which loops form and then dissociate will affect chromatin folding and this must be taken into consideration when interpreting regions of HCC.

### HCC correlation with gene expression and CTCF distribution

To begin to understand the biological role of HCC regions we have reported here that HCC correlates with increased gene expression and CTCF peak binding density. So how could HCC be associated with gene transcription? Firstly, acute loss of CTCF has been shown to alter the expression of genes within TADs in various cell types [62–64]. We speculate that the correlation we observe with CTCF may indicate that HCC may represent a local organization that enables the expression of specific genes within its local neighbourhood. In one scenario this local organisation would help bring together enhancers and promoters that need to be activated jointly. Secondly, recent microscopy based DNA tracing experiments carried out together with nascent RNA imaging have confirmed that the transcription of genes occurs in chromatin regions of highly variable folding [65–67]. Our observation that HCC regions are also highly variable from cell to cell could support a role in transcription. Thirdly, it has been suggested that the supercoiling of DNA that occurs when DNA polymerase transcribes through a gene may regulate genome folding and loop formation[68]. Thus, the HCC that we observe in discreet regions in different cells could just represent a structural feature of supercoiled chromatin that is undergoing transcription. It would now be interesting to investigate all these predictions experimentally.

## Acknowledgments

This project has received funding from the European Research Council (ERC) to D.H. under the European Union’s Horizon 2020 research and innovation programme (grant agreement No 882673). The authors acknowledge computational resources from BioClust (IBENS) facilities.

## Author contributions

A.P. and G.A. D. L. and W. B. perform research and analyze data with D. H. D.H conceive research and wrote the manuscript with the help of A.P.

## REFERENCES

[1] Lieberman-Aiden, E. et al. (2009) Comprehensive Mapping of Long-Range Interactions Reveals Folding Principles of the Human Genome. Science 326, 289–293.

[2] Rao, S. S., Huntley, M. H., Durand, N. C., Stamenova, E. K., Bochkov, I. D., Robinson, J. T., Sanborn, A. L., Machol, I., Omer, A. D., Lander, E. S., and others (2014) A 3D map of the human genome at kilobase resolution reveals principles of chromatin looping. Cell 159, 1665–1680.

[3] Cremer, T., and Cremer, C. (2001) Chromosome territories, nuclear architecture and gene regulation in mammalian cells. Nature Rev. Genet. 2, 292–301.

[4] Dixon, J. R., Selvaraj, S., Yue, F., Kim, A., Li, Y., Shen, Y., Hu, M., Liu, J. S., and Ren, B. (2012) Topological domains in mammalian genomes identified by analysis of chromatin interactions. Nature 485, 376–380.

[5] Dekker, J., Rippe, K., Dekker, M., and Kleckner, N. (2002) Capturing chromosome conformation. science 295, 1306–1311.

[6] Wang, S., Su, J.-H., Beliveau, B. J., Bintu, B., Moffitt, J. R., Wu, C.-t., and Zhuang, X. (2016) Spatial organization of chromatin domains and compartments in single chromosomes. Science 353, 598–602.

[7] Jost, D., Carrivain, P., Cavalli, G., and Vaillant, C. (2014) Modeling epigenome folding: formation and dynamics of topologically associated chromatin domains. Nucleic acids research 42, 9553–9561.

[8] Giorgetti, L., Galupa, R., Nora, E. P., Piolot, T., Lam, F., Dekker, J., Tiana, G., and Heard, E. (2014) Predictive polymer modeling reveals coupled fluctuations in chromosome conformation and transcription. Cell 157, 950–963.

[9] Brackley, C. A., Brown, J. M., Waithe, D., Babbs, C., Davies, J., Hughes, J. R., Buckle, V. J., and Marenduzzo, D. (2016) Predicting the three-dimensional folding of cis-regulatory regions in mammalian genomes using bioinformatic data and polymer models. Genome biology 17, 1–16.

[10] Amitai, A., and Holcman, D. (2017) Polymer physics of nuclear organization and function. Physics Reports 678, 1 – 83.

[11] Le Treut, G., Képès, F., and Orland, H. (2018) A polymer model for the quantitative reconstruction of chromosome architecture from HiC and GAM data. Biophysical journal 115, 2286–2294.

[12] Lando, D., Basu, S., Stevens, T. J., Riddell, A., Wohlfahrt, K. J., Cao, Y., Boucher, W., Leeb, M., Atkinson, L. P., Lee, S. F., and others (2018) Combining fluorescence imaging with Hi-C to study 3D genome architecture of the same single cell. Nature protocols 13, 1034–1061.

[13] Takizawa, T., Meaburn, K. J., and Misteli, T. (2008) The meaning of gene positioning. Cell 135, 9–13.

[14] Cremer, T., and Cremer, M. (2010) Chromosome territories. Cold Spring Harbor perspectives in biology 2, a003889.

[15] Papantonis, A., and Cook, P. R. (2013) Transcription factories: genome organization and gene regulation. Chemical reviews 113, 8683–8705.

[16] Fanucchi, S., Shibayama, Y., Burd, S., Weinberg, M. S., and Mhlanga, M. M. (2013) Chromosomal contact permits transcription between coregulated genes. Cell 155, 606–620.

[17] Zheng, H., and Xie, W. (2019) The role of 3D genome organization in development and cell differentiation. Nature Reviews Molecular Cell Biology 20, 535–550.

[18] Stevens, T. J., Lando, D., Basu, S., Atkinson, L. P., Cao, Y., Lee, S. F., Leeb, M., Wohlfahrt, K. J., Boucher, W. O?Shaughnessy-Kirwan, A., and others (2017) 3D structures of individual mammalian genomes studied by single-cell Hi-C. Nature 544, 59–64.

[19] Nagano, T., Lubling, Y., Stevens, T. J., Schoenfelder, S., Yaffe, E., Dean, W., Laue, E. D., Tanay, A., and Fraser, P. (2013) Single-cell Hi-C reveals cell-to-cell variability in chromosome structure. Nature 502, 59–64.

[20] Nagano, T., Lubling, Y., Yaffe, E., Wingett, S. W., Dean, W., Tanay, A., and Fraser, P. (2015) Single-cell Hi-C for genomewide detection of chromatin interactions that occur simultaneously in a single cell. Nature protocols 10, 1986–2003.

[21] Dekker, J., and Mirny, L. (2013) Chromosomes captured one by one. Nature 502, 45–46.

[22] Ramani, V., Deng, X., Qiu, R., Gunderson, K. L., Steemers, F. J., Disteche, C. M., Noble, W. S., Duan, Z., and Shendure, J. (2017) Massively multiplex single-cell Hi-C. Nature methods 14, 263–266.

[23] Hafner, A., and Boettiger, A. (2023) The spatial organization of transcriptional control. Nature Reviews Genetics 24, 53–68.

[24] Mach, P. et al. (2022) Cohesin and CTCF control the dynamics of chromosome folding. Nature Genetics 54, 1907–1918.

[25] Hsieh, T.-H. S., Cattoglio, C., Slobodyanyuk, E., Hansen, A. S., Darzacq, X., and Tjian, R. (2022) Enhancer–promoter interactions and transcription are largely maintained upon acute loss of CTCF, cohesin, WAPL or YY1. Nature Genetics 54, 1919–1932.

[26] Montalvão, R. W., Smith, R. E., Lovell, S. C., and Blundell, T. L. (2005) CHORAL: a differential geometry approach to the prediction of the cores of protein structures. Bioinformatics 21, 3719–3725.

[27] Leung, H., Montaño, B., Blundell, T., Vendruscolo, M., and Montalvão, R. W. (2012) ARABESQUE: A tool for protein structural comparison using differential geometry and knot theory. World Res J Peptide Protein 1, 33–40.

[28] Blundell, T., Barlow, D., Borkakoti, N., and Thornton, J. (1983) Solvent-induced distortions and the curvature of α-helices. Nature 306, 281–283.

[29] da Silva Neto, A. M., Silva, S. R., Vendruscolo, M., Camilloni, C., and Montalvão, R. W. (2019) A superposition free method for protein conformational ensemble analyses and local clustering based on a differential geometry representation of backbone. Proteins: Structure, Function, and Bioinformatics 87, 302–312.

[30] Shukron, O., and Holcman, D. (2017) Statistics of randomly cross-linked polymer models to interpret chromatin conformation capture data. Physical Review E 96, 012503.

[31] Shukron, O., and Holcman, D. (2017) Transient chromatin properties revealed by polymer models and stochastic simulations constructed from chromosomal capture data. PLoS computational biology 13, e1005469.

[32] Chang, L.-H., Ghosh, S., Papale, A., Miranda, M., Piras, V., Degrouard, J., Poncelet, M., Lecouvreur, N., Bloyer, S., Leforestier, A., and others (2021) A complex CTCF binding code defines TAD boundary structure and function. bioRxiv

[33] Leeb, M., Dietmann, S., Paramor, M., Niwa, H., and Smith, A. (2014) Genetic exploration of the exit from self-renewal using haploid embryonic stem cells. Cell stem cell 14, 385–393.

[34] Bray, N. L., Pimentel, H., Melsted, P., and Pachter, L. (2016) Near-optimal probabilistic RNA-seq quantification. Nature biotechnology 34, 525–527.

[35] Kremer, K., and Grest, G. S. (1990) Dynamics of entangled linear polymer melts: A molecular-dynamics simulation. The Journal of Chemical Physics 92, 5057–5086.

[36] Rosa, A., and Everaers, R. (2008) Structure and dynamics of interphase chromosomes. PLoS Comput Biol 4, e1000153.

[37] Bohn, M., Heermann, D. W., and van Driel, R. (2007) Random loop model for long polymers. Physical Review E 76, 051805.

[38] Bohn, M., and Heermann, D. W. (2010) Diffusion-driven looping provides a consistent framework for chromatin organization. PloS one 5, e12218.

[39] Papale, A., and Holcman, D. (2022) Chromatin phase separated nanoregions regulated by cross-linkers and explored by single particle trajectories. bioRxiv 2022–12.

[40] Alipour, E., and Marko, J. F. (2012) Self-organization of domain structures by DNA-loop-extruding enzymes. Nucleic acids research 40, 11202–11212.

[41] Fudenberg, G., Imakaev, M., Lu, C., Goloborodko, A., Abdennur, N., and Mirny, L. A. (2016) Formation of chromosomal domains by loop extrusion. Cell reports 15, 2038–2049.

[42] Polovnikov, K., Belan, S., Imakaev, M., Brandão, H. B., and Mirny, L. A. (2022) A fractal polymer with loops recapitulates key features of chromosome organization. bioRxiv

[43] Plimpton, S. (1995) Fast parallel algorithms for short-range molecular dynamics. Journal of computational physics 117, 1–19.

[44] Yang, T., Zhang, F., Yardimci, G. G., Song, F., Hardison, R. C., Noble, W. S., Yue, F., and Li, Q. (2017) HiCRep: assessing the reproducibility of Hi-C data using a stratum-adjusted correlation coefficient. Genome research 27, 1939–1949.

[45] Do Carmo, M. P. Differential geometry of curves and surfaces: revised and updated second edition; Courier Dover Publications, 2016.

[46] Crane, E., Bian, Q., McCord, R. P., Lajoie, B. R., Wheeler, B. S., Ralston, E. J., Uzawa, S., Dekker, J., and Meyer, B. J. (2015) Condensin-driven remodelling of X chromosome topology during dosage compensation. Nature 523, 240–244.

[47] Rawdon, E. J., Kern, J. C., Piatek, M., Plunkett, P., Stasiak, A., and Millett, K. C. (2008) Effect of knotting on the shape of polymers. Macromolecules 41, 8281–8287.

[48] Jagodzinski, O., Eisenriegler, E., and Kremer, K. (1992) Universal shape properties of open and closed polymer chains: Renormalization group analysis and Monte Carlo experiments. Journal de Physique I 2, 2243–2279.

[49] Hastie, T., Tibshirani, R., Friedman, J. H., and Friedman, J. H. The elements of statistical learning: data mining, inference, and prediction; Springer, 2009; Vol. 2.

[50] Nguyen, D. D., and Wei, G.-W. (2019) DG-GL: differential geometry-based geometric learning of molecular datasets. International journal for numerical methods in biomedical engineering 35, e3179.

[51] Agrawal, N. J., Radhakrishnan, R., and Purohit, P. K. (2008) Geometry of mediating protein affects the probability of loop formation in DNA. Biophysical journal 94, 3150–3158.

[52] Prior Surname, C., Davies Surname, O. R., and Pohl, E. (2019) A tertiary structure protein model for the ab-initio interpretation of small angle X-ray scattering data. bioRxiv 572057.

[53] Barbieri, M., Chotalia, M., Fraser, J., Lavitas, L.-M., Dostie, J., Pombo, A., and Nicodemi, M. (2012) Complexity of chromatin folding is captured by the strings and binders switch model. Proceedings of the National Academy of Sciences 109, 16173–16178.

[54] Shukron, O., Seeber, A., Amitai, A., and Holcman, D. (2019) Advances using single-particle trajectories to reconstruct chromatin organization and dynamics. Trends in Genetics 35, 685–705.

[55] Bianco, S., Lupiáñez, D. G., Chiariello, A. M., Annunziatella, C., Kraft, K., Schöpflin, R., Wittler, L., Andrey, G., Vingron, M., Pombo, A., and others (2018) Polymer physics predicts the effects of structural variants on chromatin architecture. Nature genetics 50, 662–667.

[56] Shi, G., and Thirumalai, D. (2023) A maximum-entropy model to predict 3D structural ensembles of chromatin from pairwise distances with applications to interphase chromosomes and structural variants. Nature Communications 14, 1150.

[57] Takaki, R., Dey, A., Shi, G., and Thirumalai, D. (2021) Theory and simulations of condensin mediated loop extrusion in DNA. Nature Communications 12, 5865.

[58] Fiorillo, L., Musella, F., Conte, M., Kempfer, R., Chiariello, A. M., Bianco, S., Kukalev, A., Irastorza-Azcarate, I., Esposito, A., Abraham, A., and others (2021) Comparison of the Hi-C, GAM and SPRITE methods using polymer models of chromatin. Nature Methods 18, 482–490.

[59] Sanborn, A. L., Rao, S. S., Huang, S.-C., Durand, N. C., Huntley, M. H., Jewett, A. I., Bochkov, I. D., Chinnappan, D., Cutkosky, A., Li, J., and others (2015) Chromatin extrusion explains key features of loop and domain formation in wild-type and engineered genomes. Proceedings of the National Academy of Sciences 112, E6456–E6465.

[60] Gabriele, M., Brandão, H. B., Grosse-Holz, S., Jha, A., Dailey, G. M., Cattoglio, C., Hsieh, T.-H. S., Mirny, L., Zechner, C., and Hansen, A. S. (2022) Dynamics of CTCF-and cohesin-mediated chromatin looping revealed by live-cell imaging. Science 376, 496–501.

[61] Hansen, A. S., Pustova, I., Cattoglio, C., Tjian, R., and Darzacq, X. (2017) CTCF and cohesin regulate chromatin loop stability with distinct dynamics. elife 6, e25776.

[62] Nora, E. P., Goloborodko, A., Valton, A.-L., Gibcus, J. H., Uebersohn, A., Abdennur, N., Dekker, J., Mirny, L. A., and Bruneau, B. G. (2017) Targeted degradation of CTCF decouples local insulation of chromosome domains from genomic compartmentalization. Cell 169, 930–944.

[63] Kubo, N., Ishii, H., Xiong, X., Bianco, S., Meitinger, F., Hu, R., Hocker, J. D., Conte, M., Gorkin, D., Yu, M., and others (2021) Promoter-proximal CTCF binding promotes distal enhancer-dependent gene activation. Nature structural & molecular biology 28, 152–161.

[64] Stik, G., Vidal, E., Barrero, M., Cuartero, S., Vila-Casadesús, M., Mendieta-Esteban, J., Tian, T. V., Choi, J., Berenguer, C., Abad, A., and others (2020) CTCF is dispensable for immune cell transdifferentiation but facilitates an acute inflammatory response. Nature genetics 52, 655–661.

[65] Mateo, L. J., Murphy, S. E., Hafner, A., Cinquini, I. S., Walker, C. A., and Boettiger, A. N. (2019) Visualizing DNA folding and RNA in embryos at single-cell resolution. Nature 568, 49–54.

[66] Gizzi, A. M. C., Cattoni, D. I., Fiche, J.-B., Espinola, S. M., Gurgo, J., Messina, O., Houbron, C., Ogiyama, Y., Papadopoulos, G. L., Cavalli, G., and others (2019) Microscopy-based chromosome conformation capture enables simultaneous visualization of genome organization and transcription in intact organisms. Molecular cell 74, 212–222.

[67] Takei, Y., Zheng, S., Yun, J., Shah, S., Pierson, N., White, J., Schindler, S., Tischbirek, C. H., Yuan, G.-C., and Cai, L. (2021) Single-cell nuclear architecture across cell types in the mouse brain. Science 374, 586–594.

[68] Neguembor, M. V., Martin, L., Castells-García, Á., Gómez-García, P. A., Vicario, C., Carnevali, D., Abed, J. A., Granados, A., Sebastian-Perez, R., Sottile, F., and others (2021) Transcription-mediated supercoiling regulates genome folding and loop formation. Molecular cell 81, 3065–3081.

